# Uromonitor^®^ as a novel sensitive and specific urine-based test for recurrence surveillance of patients with non-muscle invasive bladder cancer

**DOI:** 10.1101/410738

**Authors:** Cristina Sampaio, Rui Batista, Pedro Peralta, Paulo Conceição, Amílcar Sismeiro, Hugo Prazeres, João Vinagre, Paula Soares

## Abstract

Bladder cancer is the most frequent malignancy of the urinary system and is ranked the seventh most diagnosed cancer in men worldwide. About 70-75% of all newly diagnosed patients with bladder cancer will present disease confined to the mucosa or submucosa, the non-muscle invasive bladder cancer (NMIBC) subtype. Of those, approximately 70% will recur after transurethral resection (TUR). Due to this high rate of recurrence, patients are submitted to an intensive follow-up program that should be maintained throughout many years, or even throughout life, resulting in an expensive follow-up, with cystoscopy being the most cost-effective procedure for NMIBC screening. Currently, the gold standard procedure for detection and follow-up of NMIBC is based on the association of cystoscopy and urine cytology. As cystoscopy is a very invasive approach, over the years, many different non-invasive (both in serum and urine samples) assays have been developed in order to search genetic and protein alterations related to the development, progression and recurrence of bladder cancer. *TERT* promoter mutations and *FGFR3* hotspot mutations are the most frequent somatic alterations in bladder cancer and constitute the most reliable biomarkers for bladder cancer. Based on these findings, an ultra-sensitive assay called Uromonitor^®^ was developed that corresponds to a urine-based assay capable of detecting trace amounts of the two most common alterations in NMIBC, *TERT* promoter and *FGFR3* mutation, in urine samples. The Uromonitor^®^ test was performed in a cohort of 72 patients, firstly diagnosed with bladder cancer and under surveillance for NMIBC, to access its sensitivity and specificity in the detection of NMIBC recurrence. Uromonitor^®^ was shown to be highly sensitive and specific in detecting recurrence of bladder cancer in patients under surveillance of non-muscle invasive bladder cancer.

## Introduction

Bladder cancer is the most frequent malignancy involving the urinary system and affects approximately 4 times more the male gender than the female [1]. Worldwide, bladder cancer is the seventh cancer most diagnosed in men; when both genders are considered it ranks the eleventh position [2]. Of all newly patients diagnosed with bladder cancer, around three quarters present disease confined to the mucosa or submucosa [3], the non-muscle invasive bladder cancer (NMIBC) subtype [4]. The remaining are classified as muscle invasive bladder cancer (MIBC) reflecting their capacity to infiltrate the muscle layer of the bladder [3, 5]. The gold standard for NIMBC treatment is the transurethral resection (TUR); following TUR treatment, 70% of the NMIBC patients will recur after primary tumor removal and 10 to 20% will recur as muscle-invasive bladder cancer and with the capacity to progress and develop metastatic disease [6-8]. This high rate of recurrence requires that patients are submitted to an intensive follow-up program. Major guidelines from European Association of Urology (EAU) and Canadian Urological Association (CUA) recommend cystoscopy and urinary cytology every 3 months in the first 2 years, semi annually during the subsequent 3 years and annually thereafter [9]. This intensive follow-up is maintained throughout many years following the initial diagnosis and confers bladder cancer as a type of cancer with the most expensive follow-up; consequently, cystoscopy is the most cost-effective procedure for follow-up of NMIBC [10, 11]. Currently, the gold standard to detect bladder cancer recurrence is based on the association of cystoscopy and urine cytology [4, 5]. Cystoscopy is invasive and uncomfortable for patients due to the technical requirements of the procedure, still, it renders the more accurate diagnosis method for bladder cancer [12]. Contrarily to cystoscopy, the non-invasive urine cytology is an economical approach, easier to perform, and when high-grade tumors are considered, the sensitivity is high (84%). The major limitation of urine cytology is its overall sensitivity to detect tumor cells that decreases to 16% in low-grade tumors, precluding its use in detection of those lesions [13]. The combination of all these facts leads to the opportunity for developing new, alternative and minimally invasive methods to detect bladder cancer. As urine is in direct contact with the inner part of bladder, cells from the epithelium, including scammed cells from bladder tumors can exfoliate and be detected in urine and used to evaluate and monitor the presence of neoplasia in a non-invasive approach [14-18]. Over the years, many different non-invasive assays have been developed in order to search genetic and protein alterations well known to be involved in the development, progression and recurrence of bladder cancer, both in serum and urine samples with the purpose to diagnose and monitor bladder cancer [13, 14, 19-35]. Some of these tests present values of sensitivity and specificity higher than urinary cytology and achieved FDA-approval for bladder cancer diagnosis. Despite high sensitivities and specificities, all these molecular assays present inconvenient rates of false positive results [34, 36-38]. False positive rates could result from several factors, including the presence of benign conditions as hematuria, cystitis, lithiasis, urinary tract infections, inflammation or even because of repeated instrumentation, such as cystoscopy [39, 40]. A meta-analysis about the performance of urinary biomarkers conclude that most of the available urinary biomarkers do not detect the presence of bladder cancer in a proportion of patients and allow false-positive results in others, more frequently in low-stage and low-grade tumors [41]. Despite their good performance when combined with each other or even with urinary cytology, more reliable biomarkers and assays are needed for earlier detection of bladder cancer recurrence, particularly in low-grade and low-stage NMIBC. Recently, telomerase reverse transcriptase (TERT) promoter mutations emerged as a novel biomarker and detected in up to 80% of bladder cancer, independently of stage or grade [30, 31, 42, 43]. *TERT* promoter mutations are the most common event across stages and grades in malignant bladder tumors, strongly suggesting its participation in the two major genetic pathways of urothelial tumorigenesis [30, 31]. These features point *TERT* promoter mutations to be considered as a useful urinary biomarker for disease monitoring and early detection of recurrence, even in low-grade NMIBC, where urinary cytology usually lacks sensitivity [30, 31, 44, 45]. *TERT* promoter mutations are not present in inflammatory or urinary infections, different from previously described urinary biomarkers [41, 45, 46]. *TERT* promoter mutations assumed a novel pivotal role, surpassing the frequency of the oncogene-activating mutations in fibroblast growth factor receptor 3 (FGFR3) gene in NMIBC [47, 48]. Cappellen *et al*., reported *FGFR3* mutations in bladder cancer with a frequency of 35% and subsequent studies established this frequency in approximately half of the primary bladder tumors [49, 50]; several studies report its presence in up to 80% regarding early stage and low-grade tumors, and as absent or a very rare event in high-grade and invasive tumors [51-55].

Based on these findings, we developed an ultra-sensitive assay named Uromonitor^®^, a urine-based test capable of detecting trace amounts of the most common alterations in NMIBC, *TERT* promoter and *FGFR3* hotspot mutations, in urine samples.

## Material and Methods

### Sample collection

#### Urine samples

Urine samples were collected during routine urology appointments and previously to cystoscopic intervention. The samples were centrifuged in a 50 mL centrifuge tube at 300 g for 20 minutes at room temperature (15 - 30°C). Supernatant was carefully removed and pellets were stored at -80^ª^C until DNA extraction procedure.

#### Tissue samples

FFPE tissues from primary tumor and/or from recurrent lesions from the cohort in study were retrieved from the files of the Instituto Português de Oncologia de Coimbra Francisco Gentil, E.P.E (IPOC-FG). Clinicopathological and follow-up data were retrieved from the files of the Department of Pathology of IPOC-FG. For the technical validation we used also a series of thyroid tumors FFPE samples (n=196) retrieved from the files of IPATIMUP.

### Cohort characteristics

#### Clinical validation

We studied a total of 139 samples corresponding to urine (n=98) and formalin-fixed paraffin-embedded (FFPE) tissues of the primary tumor and of the recurrence lesions (n=41) from a cohort of patients under follow-up after an initial diagnosis and treatment for bladder cancer. When available, FFPE tissues of the corresponding primary tumor (n=9) and/or of the recurrence lesions (n=32) were also analyzed. A case was considered positive for recurrence when the cystoscopy evaluation reported the possibility of recurrence, and malignancy was confirmed in the histological examination of the recurrence lesion in the TUR. All procedures described in this study were in accordance with national and institutional ethical standards and previously approved by Local Ethical Review Committees.

#### Technical validation

We studied a total of 334 samples corresponding to urine (n=97) and formalin-fixed paraffin-embedded (FFPE) tissues (n=237) derived from bladder cancer tumor samples, bladder inflammatory tissue and thyroid tumors (only FFPE samples). From the 97 urine samples, 73 were successfully analysed for TERT -124 assay, 72 for TERT -146 assay, 55 for FGFR3 248 assay and 52 for FGFR3 249 assay. From the 237 FFPE tissue samples, 201 were successfully analysed for TERT -124 assay, 200 for TERT -146 assay, and 41 for both FGFR3 248 and FGFR3 249 assays. All procedures described in this study were in accordance with national and institutional ethical standards and previously approved by Local Ethical Review Committees.

### DNA extraction

#### Urine samples

Pellets stored at -80°C were thawed at room temperature. Pellets of cells were resuspended in 25 mL of PBS 1x (Phosphate-buffered saline solution) and filtered in nitrocellulose membranes to retain cells. Filters were put on top of a 2 mL microcentrifuge tube in the reverse position of filtering and a cell lysis solution was added to each filter and passed through to the tube. Filtered lysates were incubated at 60°C for 10 min with 30μL of Proteinase K and exposed to chaotropic lysis/binding buffer to release nucleic acids and protect the genomic DNA from DNases. The microcentrifuge tubes content was then processed according to the manufacturer’s protocol of the Norgen® Plasma/Serum Cell-Free Circulating DNA Purification Mini Kit (Norgen Biotek Corp, Canada).

#### Tissue samples

DNA from FFPE tissues was retrieved from 10-μm cuts after careful manual dissection. Slides were deparaffinized in xylene (2× 10 minutes), followed by incubation in 100% alcohol (2× 5 minutes). All tumor tissue was removed from the slides to a 1.5 mL microcentrifuge tube. DNA extraction was performed using the Ultraprep Tissue DNA Kit (AHN Biotechnologie, Germany) following the manufacturer’s instructions.

### DNA quantification

DNA extracted from both sample types was quantified spectrophotometrically using Nanodrop ND-2000, and its quality was checked by analysis of 260/280nm and 260/230 nm ratios.

### Uromonitor^®^ test

The Uromonitor® test was developed and optimized in a custom made full working procedure for the detection of TERT and FGFR3 hostpot mutations in bladder cancer tumor cells exfoliated to urine.

Mutation detection is achieved by Real-Time PCR amplification curve analysis. Positive and negative mutation control samples are included in each run to ensure assay’s validity. For each of the selected mutations in screening, we designed a mutant allele-specific primer (ASP), a wild type allele-specific blocker (ASB), a common reverse primer (LSP) for both alterations and a TaqMan probe for Real-Time detection of the generated amplicon. The use of a molecular blocker completely suppresses the amplification of the wild type allele to not interfere with the amplification of the mutant allele. By this technique we improved current detection thresholds enhancing the ability to detect a minimal quantity of altered cells in a large pool of cell without alterations.

Uromonitor^®^ test was performed in approximately 25 ng of DNA extracted from cells in each filtered urine, or from 25 ng of DNA extracted from FFPE tissues, either primary tumor and/or recurrent lesions. The extracted DNA was amplified and detected on a qPCR real-time machine, using the proprietary chemistry for amplification and detection as provided in the Uromonitor^®^ test kit for the targeted nucleotide changes in *TERT* promoter and *FGFR3*.

### Uromonitor^®^ technical validation

Uromonitor’s precision was analysed by a reproducibility test. To ensure this, 10 samples were amplified and analysed using Uromonitor® test (8 samples harbouring mutations and 2 samples wildtype for the alterations of interest). This samples were amplified 5 times for each alteration, 1 week apart of each amplification, during 5 weeks. Uromonitor’s accuracy was analysed by two independent tests. First, it was necessary to ensure that a test containing a sample without DNA or with DNA that doesn’t harbour any of the alterations of interest, do not generate an analytical signal that may indicate a low concentration of mutation (false positive). It was also necessary to access the accuracy of the results produced by Uromonitor® test comparing it to the gold standard method in the detection of the alterations in study (Sanger Sequencing). To achieve this samples amplified by Sanger Sequencing for the alterations in study were also amplified by Uromonitor® test.

### Uromonitor’s quality assessment by the determination of precision and accuracy of the obtained results in urine samples

To test accuracy in urine samples, 36 samples negative for all the mutations in study (status obtained by Sanger Sequencing) and 36 “Blank” samples (without DNA) were amplified for each alteration (false-positive testing).

Also, 252 blind tests from urine samples were analysed (73 tests for TERT -124 assay, 72 tests for TERT -146 assay, 55 tests for FGFR3 248 assay and 52 tests for FGFR3 249 assay).

### Uromonitor’s quality assessment by the determination of precision and accuracy of the obtained results in FFPE tissue samples

Uromonitor’s test could also be used to screen initial tumor from patients with history of NMIBC. With this the eligibility of the patient to perform the test with full confidence is assured. Uromonitor’s test was also optimized to work with FFPE tissue.

To test accuracy in FFPE tissue samples, 9 samples negative for all the mutations in study (status obtained by Sanger Sequencing), 36 “Blank” samples (without DNA) were amplified for each alteration (false-positive testing). Also, 483 tests from FFPE tissue samples were analysed (201 tests for TERT -124 assay, 200 tests for TERT -146 assay, 41 tests for FGFR3 248 assay and 41 tests for FGFR3 249 assay). In this test, to increase the number of samples in the analysis of TERTp, we included also 196 FFPE thyroid cancer samples previously analysed by Sanger sequencing.

### Detection limit threshold

Detection limit threshold was obtained for both TERTp and FGFR3 alterations that compose the Uromonitor®. To achieve this, we performed 2-fold serial dilutions of genomic DNA containing the studied alteration (100% of mutated DNA) in genomic DNA wildtype for the studied alterations. Serial dilutions were amplified for the corresponding detection assay. This procedure was repeated for all the studied alterations that comprise the Uromonitor® test.

## Results

### Technical validation

Uromonitor’s precision was analysed, achieving 100% concordance in a reproducibility test.

Regarding Uromonitor’s test accuracy in urine samples (comparison to Sanger sequencing), TERT -124 assay achieved 100%, TERT -146 assay 98,6%, FGFR3 248 assay 87,3% and FGFR3 249 assay 94,2%. Overall, Uromonitor® test achieved a combined 95% accuracy. Regarding Uromonitor’s test accuracy in FFPE tissue samples (comparison to Sanger Sequencing), TERT -124 assay achieved 98,5%, TERT -146 assay 99,5%, FGFR3 248 assay 90,2% and FGFR3 249 assay 97,6%. Overall, Uromonitor® test achieved a combined 96,5% accuracy, Table 1. Combined accuracy is lower than 100%, and it could be partially justified by the detection of positivity in samples in which Sanger sequencing failed to detect alteration due to lack of sensitivity. Also, the test presented no false positives in samples without DNA (Blank samples).

**Table 1:** Uromonitor^®^ technical validation – accuracy

|  | **-124 Assay** | **-146 Assay** | **FGFR3 248** | **FGFR3 249** | **Combined Assays** |
| --- | --- | --- | --- | --- | --- |
| **Urine samples** | 100% | 98.6% | 87.3% | 94.2% | 95.0% |
| **FFPE samples** | 98.5% | 99.5% | 90.2% | 97.6% | 96.5% |

For all the assays, the limit threshold was defined at ≥ 6.25% mutant sequences in a background of wild-type DNA. This means that in the presence of a DNA sample in which altered DNA presents less than 6.25% of total DNA present in the sample, it is not granted that the assay could detect the alteration.

### Clinical validation

From an initial cohort of 98 urine samples, 72 patients were fully characterized for the alterations targeted by Uromonitor^®^ test. In this cohort, 22.9% of cases presented recurrence, as reported by the cystoscopy examination and confirmed by histology of the TUR biopsy(ies), whereas the remaining 77.1% were negative for recurrence, Table 2. Of note, 5.6% of cystoscopic examinations in our series reported a recurrence, yet the histological examination of the subsequent TUR biopsies yielded only benign and/or inflammatory changes, and no recurrence was detected in the two years of follow-up of these patients, which can be taken to indicate that these are cystoscopy false positives. All cases with a positive recurrence were positive for at least one or more of the mutations detected by the Uromonitor^®^ test, resulting in a sensitivity of 100%. It is noteworthy that in recurrence positive cases (by cystoscopy and histology) in which benign/inflammatory lesions coexisted with carcinoma lesions (comprising 23% of the positive recurrences) genetic analysis of the different lesions/components revealed that mutations were exclusively detected in the malignant counterparts and were not detected in the benign lesion/component. Concomitant urine cytology performed in the recurrence positive cases was compatible with recurrence or suspected of malignancy in only 38.0% of cases, with 42.0% of cases having a report of absence of neoplastic cells and/or inflammation and the remaining 15.0% were reported as inconclusive.

**Table 2:** Correlation between Mutation Status in Urine and Recurrence

| **Mutation Status**<br>**in Urine Samples** | **Recurrence** |  |
| --- | --- | --- |
|  | **Positive**<br>*n*=16<br>(22.9%) | **Negative**<br>*n*=52<br>(77.1%) |
| **Positive** *n*=18, (25,7%) | 16 (22.9%) | 2 (2.9%) |
| **Negative** *n*=52, (74,3%) | 0% | 52 (74.3%) |

Regarding the specific mutations detected in recurrence positive cases, *TERT* promoter mutations were detected in 53% of cases (42% presented the -124C>T and 11% with the -146C>T) and *FGFR3* mutations were detected in 47% of cases (32% at codon 248 and 16 at codon 249 of FGFR3 protein).

Amongst the recurrence negative cases, the Uromonitor^®^ test was concordantly negative for mutations in 96.3% of cases, with the remaining 2.9% (2 cases) presenting mutations for *FGFR3* in codon 248, despite negativity for recurrence. These cases have remained negative for recurrence in the 2 years follow-up, suggesting these were false positives. On the basis of these findings, the test sensitivity was 100% and specificity was 96.3%. The negative predictive value was 100%, and the positive predictive value was 88.9%, Table 3.

**Table 3:** Uromonitor^®^ assay biological validation

|  |  |
| --- | --- |
| **Sensitivity** (*mut+/rec+*) | 100% |
| **Specificity** (*mut-/rec-*) | 96.3% |
| **Positive predictive value** | 100% |
| **Negative predictive value** | 88.9% |

*mut*: mutation; *rec*: recurrence

Since test sensitivity was 100%, all recurrences were detected, irrespective of their stage and grade. The majority of recurrence positive cases were in stage Ta (46.0%), with T1 and Tis representing 23.0% and 31.0%, respectively. Regarding the grade, the majority of recurrence positive tumours were low grade (69.0%) being the remaining (31.0%) high grade cases.

Comparison of the genetic profiles between urine and recurrence lesion of the corresponding patient, 60% of the cases hold at least one of the mutations detected in urine that was also identified in the corresponding recurrence lesion, with two thirds of these cases showing additional mutations in the recurrence lesion analysis. The remaining 40% of cases rendered distinct mutations in the urine and recurrence lesion.

## Discussion

*TERT* promoter mutations were firstly described in sporadic and familial melanoma [58, 59] and since then they were reported in several cancers, such as central nervous system (43-51%), hepatocellular carcinoma (59%), thyroid (follicular cell-derived tumors) (10%) and notably in bladder cancer (59-80%) [60-65]. For bladder cancer, the *TERT* promoter mutations are independent of stage or grade [30, 31, 42, 43] and were reported in both non-muscle and muscle invasive bladder cancer. As the current treatment of patients with bladder cancer is based on a very variable analysis from pathologists, based mainly on stage and grade of the tumor, new molecular markers are needed to better characterize and classify bladder cancer patients and to select an optimal treatment for each [55, 66, 67]. For this purpose, we developed a novel real-time assay, Uromonitor^®^, and in this study we assess the technical and clinical performance of the detection of alterations in *TERT* promoter region, and on *FGFR3* gene in DNA obtained from scammed cells of bladder present in urine. The main goal of Uromonitor^®^ is to be able to predict recurrence in NMIBC. The initial cohort was composed of 98 urine samples, however, due to material quality, only 72 urine sample from patients were able to be fully characterized for the alterations targeted by Uromonitor^®^ test. This was a consequence of the sample preparation procedures adopted (inefficient freezing procedures and longtime storage); samples processed and analyzed following Uromonitor^®^ test collection procedures do not present this pitfall.

The Uromonitor^®^ test presented a 100% specificity in the detection of recurrence and with equally performance within different grade and stage. For the cases with different components of benign and tumour counterparts the analysis was able to discern the malignant ones. These findings indicate that Uromonitor^®^ test can be put into practice as an aid for clinicians to distinguish recurrence from inflammatory tissue after a suspicious or abnormal cystoscopy [45]. In this series, recurrence was detected (by histologic confirmation) in 22.9% of the studied cases, a value distant from the 60 to 70% reported in the literature for NMINBC following TUR treatment [6-8]. At this moment, it remains to be elucidated the reason for this difference in the recurrence values, but we can assume that they reflect patient-related factors (age, gender, multiplicity, smoking status and adjuvant treatment) associated with recurrence frequency [68] or the restricted two year follow-up that patients were considered in this study.

The mutational state of *FGFR3* oncogene in bladder cancer is considered a promising predictor for recurrence and progression of NMIBC demonstrated by the association between *FGFR3* mutation in primary tumor and later in recurrence events [53, 69-73]. In line with previous evidence *FGFR3* was detected more frequently in recurrent cases and surpassing the most common genetic event in bladder cancer the *TERT* promoter mutations. Still, two cases were positive for *FGFR3* mutations without evidence for recurrence after the follow-up terminus that dropped the specificity of Uromonitor^®^ to 96.3%. More than half of the cases presented at least one mutational event and it is reported that *TERT* promoter and *FGFR3* mutations tend to occur more frequently together than per chance; the combination of both might constitute a more reliable biomarker for NMIBC recurrence monitoring [15, 42]. In terms of the Uromonitor^®^ performance in comparison with other available options it presented improved features. Reviewed by Sapre *et al.* [74] the sensitivity of other available options ranges from 50.0% to 96.6%, and the tests are based in different methodologies approaches, some more technically challenging and maintaining invasive requirements for the procedure in the patients. Avoidance of invasive procedures for the patients was a concern in the test development since morbidity of cystoscopy is often underestimated and can impact on patient adherence with surveillance as low as 40% [75]. The fact that the test is conducted in urine, renders it safer for patient use and with better acceptance in comparison with conventional cystoscopy.

Overall, our study demonstrates that Uromonitor^®^ represents a highly sensitive and specific urine test in detecting recurrence of NMIBC, and this outstanding sensitivity was maintained in the different tumor stages and/or grades. Due to its extremely high negative predictive value, the test could potentially alternate with cystoscopy in surveillance programs, diminishing the risk of missing recurrences, and with the benefit of alleviating the number cystoscopy procedures that patients require to undergo. The rate of Uromonitor^®^ false positives was actually lower than the rate of cystoscopy false positives. Our results prompt us to validate these findings in an independent series, in an ongoing study with a design that includes a group of benign conditions (renal lithiasis, urinary infections, hyperplasia of the prostate or other) and a head-to-head comparison with cytology. We intent to further test it to assess its cost-effectiveness and to determine its value in patients follow-up.

